# The shapes of wine and table grape leaves: an ampelometric study inspired by the methods of Pierre Galet

**DOI:** 10.1101/2020.05.08.085175

**Authors:** Daniel H. Chitwood

**Author notes:** To whom correspondence should be addressed: Daniel H. Chitwood, Michigan State University, Dept. Horticulture, 1066 Bogue St., East Lansing, MI 48824 USA.

## Abstract

The shapes of grapevine leaves have been critical to correctly identify economically important varieties throughout history. The correspondence of homologous features in nearly all grapevine species and varieties has enabled advanced morphometric approaches to mathematically classify leaf shape. These approaches either model leaves through the measurement of numerous vein lengths and angles or measure a finite number of corresponding landmarks and use Procrustean approaches to superimpose points and perform statistical analyses. Hand illustrations, too, play an important role in grapevine identification, as details omitted using the above methods can be visualized. Here, I use a saturating number of pseudo-landmarks to capture intricate, local features in grapevine leaves: the curvature of veins and the shapes of serrations. Using these points, averaged leaf shapes for 60 varieties of wine and table grapes are calculated that preserve features. A pairwise Procrustes distance matrix of the overall morphological similarity of each variety to the other classifies leaves into two main groups—deeply lobed and more entire—that correspond to the measurements of sinus depth by Pierre Galet. Using the system of Galet, pseudo-landmarks are converted into relative distance and angle measurements. Both Galet-inspired and Procrustean methods allow increased accuracy in predicting variety compared to a finite number of landmarks. Using Procrustean pseudo-landmarks captures grapevine leaf shape at the same level of detail as drawings and provides a quantitative method to arrive at mean leaf shapes representing varieties that can be used within a predictive statistical framework.

## INTRODUCTION

The grapevine leaf is a coordinate system defined by vasculature, the branching points and termination of which can be found in nearly all *Vitis* spp. leaves. Each leaf has a midvein, two distal/superior veins, two proximal/inferior veins, and two prominent veins that branch off of the proximal veins called petiolar veins (**Fig. 1**). The major primary veins of the leaf terminate at the lobe tips. The secondary veins that branch off the primary terminate at the blade margin, forming serration patterns between consecutive branches. Using the ordered branching pattern that emerges from the primary veins defining each lobe, a hierarchy of venation and serrated teeth along the blade can be defined. This system permits spatial correspondence between all grapevine leaves that has enabled sophisticated morphometric approaches and historical application to the discrimination of species and varieties using leaf shape.

**Figure 1:**
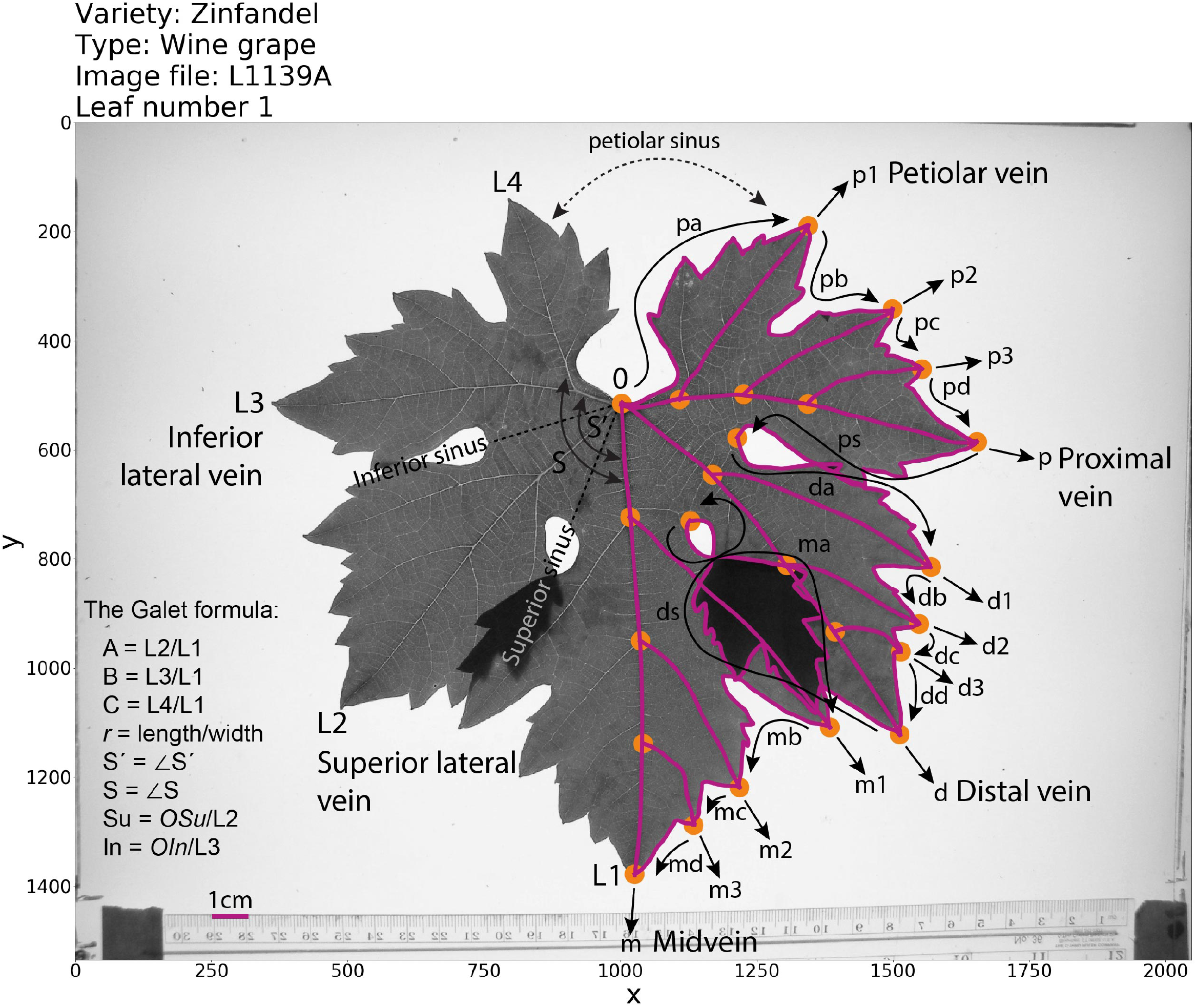
The Galet formula and Procrustean methods. A scan of a Zinfandel leaf over which raw data has been plotted. Data is saved as image pixel coordinates. On the right side of the leaf landmarks (orange dots) and pseudo-landmarks (magenta lines) are plotted. Landmark data are saved as vectors, the names of which are indicated next to corresponding arrows. “p”, “d”, and “m” refer to “proximal”, “distal”, and “midvein” regions of the leaf. Along the blade, the base of each arrow and its tip indicate the beginning and end of a vector. Arrows arising from the tips of veins indicate the direction of vein vectors that originate at corresponding branch points within the leaf and terminate at the tips. On the left side of the leaf, the nomenclature of Galet is provided. Midvein, distal/superior, proximal/inferior, and petiolar veins are called L1, L2, L3, and L4, respectively. Superior and inferior sinuses are shown, as well as angles S’ and S between L1/L3 and L1/L4, respectively. A, B, and C are ratios of the lengths of L2, L3, and L4, respectively, to L1; *r* is the ratio of length to width; and Su and In are the distances to the petiolar junction (0) of the superior (Su) and inferior (In) sinuses divided by the length of the L2 and L3, respectively.

In the mid-1850s, an aphid crossed the Atlantic from North America attacking the root system of *Vitis vinifera* (domesticated grape) vines in France decimating the wine industry. North American *Vitis* spp. rootstocks were resistant to the pest and ultimately the solution to the blight that restored wine production. The rootstocks were new to European viticulturists, yet correctly identifying and selecting the correct rootstock variety was vital. The roots themselves and the grape clusters were of little use to identifying varieties, so viticulturists turned to the leaves. The field of ampelography (“vine” + “writing”), concerning the discrimination of grapevine varieties, was born and chief among its techniques was ampelometry (“vine” + “process of measuring”), a method of measuring leaf shape. Hermann Goethe (Goethe, 1876; 1878) first proposed to use the angle of the petiolar sinus to identify North American *Vitis* spp., but Louis Ravaz expanded upon the idea and established a foundational system for quantifying the shapes of grapevine leaves in his *Les vignes américaines: Porte-greffes et producteurs directs* (1902). A focus on not only the angle, but the shape and contour of the petiolar sinus in hand-drawings was made. The overall shape of the leaf (reniform, orbicular, cordiform, cuneiform, or truncate) was described in terms of ratios of lengths and angles between veins, and codified into discrete groups based on ranges of values. Even the serrations were described in terms of length-to-width ratio and convex/concave shapes.

While Ravaz popularized the system of ampelometry, Pierre Galet turned it into an artform (Galet 1979; 1985; 1988; 1990; 2000). In his works, Galet hand draws a representative leaf for each variety, a format that guides the reader’s eyes to the major veins and their relationship to the blade. Extensive information regarding the history, geography, and phenology of vines, and the appearance of the inflorescence and growing tip, in addition to descriptions of leaf hirsuteness, contour, and surface, verbally recreates the experience of encountering a vine in the reader’s mind. Like Ravaz, Galet created a discretized system of values to describe ratios of vein lengths and angles (the Galet formula), but also created measuring devices (the Galet ruler and protractor) to easily quantify values in the vineyard and compare to ideal values for each variety that he published. Galet, through careful observation, a quantitative mindset, detailed description, encyclopedic knowledge, hand illustration, and an artist’s eye effectively transcribed the immense phenotypic variation among *Vitis* spp. into books that have since inspired and taught those who work with and love grapevines.

Others took the analysis of grapevine leaves in a more mathematical direction. The homologous coordinates in every *Vitis* spp. leaf allows even minor veins to be hierarchically accounted for. By counting teeth, where veins terminate, and measuring leaf shape, Acúrcio Rodrigues developed a method for calculating an average leaf shape (Rodrigues 1939; 1941a; 1941b; 1952a; 1952b). María-Carmen Martínez developed the method further, and through statistically measuring numerous angles, lengths, and numbers of teeth for a variety, developed a model for reconstituting a visual representation of an average leaf (Martínez and Grenan, 1999). The method opened the door for statistical analysis of grapevine leaf morphology (Martinez et al., 1995), discriminating cultivars (Santiago et al., 2005; 2007; 2008; Gago et al., 2009a), clones (Martínez et al., 1997a; 1997b), and even comparing depictions of leaves in historical works of art to present-day varieties (Gago et al., 2009b; 2014).

Another approach to measuring shape is landmarks (Booketein, 1997): homologous *x* and *y* coordinates that are found in every leaf. Using Procrustean methods, landmarks can be superimposed through translation, rotation, scaling, and reflection minimizing the distance of all points to each other (Gower, 1975). Although landmarks capture less of the overall shape of an object, because they are finite, high levels of replication are possible. Tens of thousands of grapevine leaves have been measured using landmarks. Previous analysis of wine and table grape varieties in the USDA Wolfskill National Clonal Germplasm Repository in Winters, California (USA) used ten landmarks along the distal and proximal lobe tips and sinuses (excluding the petiolar veins) on both sides of the leaf to measure the genetic basis of leaf shape (Chitwood et al., 2014). A set of 17 landmarks including the petiolar vein and the first major secondary branch points of the midvein, distal vein, and proximal vein on both sides of the leaf was used to explore leaf shape in a developmental and evolutionary context using *Vitis* spp. in the USDA Geneva, New York (USA) germplasm collection (Chitwood et al., 2016a), to find conserved loci regulating leaf shape in multiple *Vitis* spp. interspecific hybrid mapping families (Demmings et al., 2019), and to document inter- and intra-species leaf shape variation between *V. riparia* and *V. rupestris* clones at the Missouri Botanical Garden, St. Louis (USA; Klein et al., 2017). A set of 21 landmarks capturing the widths of the primary veins and their major secondary branching veins for half of the leaf was used to reanalyze the USDA Geneva, New York (USA) germplasm across two years on the same vines to test for climate-induced changes in leaf shape plasticity (Chitwood et al., 2016b).

Although insightful and permitting the analysis of thousands of leaves, a finite number of landmarks fails to capture the curves, serrations, and intricate details of grapevine leaf shape that are readily apparent by eye. The analysis presented here attempts to capture these finer features of grapevine leaf shape by 1) maximizing the number of landmarks used and 2) capturing curves and local features (such as serrations) by using a saturating number of pseudo-landmarks between them.

## MATERIALS AND METHODS

### Plant material and photography

Over 9,500 leaves from more than 1,200 wine and table grape varieties *(Vitis vinifera)* were collected at the USDA Wolfskill National Clonal Germplam Repository in Winters, California (USA) from May 28 through June 1, 2011. As previously described in Chitwood et al., 2014, photographs of the leaves were taken using a remote-controlled camera attached to a copy stand and placing the leaves under nonreflective glass to flatten them on top of a light box to highlight venation. A total of 4,950 photos were taken, named by vine location that serves as a key for variety identity. In the previous study, the shapes of all leaves were measured using ten landmarks. This study examines a small subset of 60 varieties in intensive detail that were also described by Pierre Galet in *A Practical Ampelography* (Galet, 1979; 1985). The original photographs used for this study can be found at https://github.com/DanChitwood/grapevine_ampelometry/tree/master/0_visual_check/ampel_ometry_images. Each photo is named by its vineyard location at the USDA Wolfskill repository followed by letters if multiple images were taken for the sampled clones, which can be used to determine variety identity using the following key: https://github.com/DanChitwood/grapevine_ampelometry/blob/master/0_visual_check/ampel_ometry_id_key.txt

Only 60 varieties are analyzed in this study, but all of the >4,950 photos of 9,500 leaves of more than 1,200 wine and table grape varieties can be downloaded using the following doi at Dryad: https://doi.org/10.5061/dryad.g79cnp5mn

### Landmarking, tracing, and visual checks

24 landmarks corresponding to the tips of midvein, distal vein, and proximal vein (3 points), the distal and proximal sinuses (2 points), the petiolar junction (1 point), and the three major secondary branch points for the midvein, distal vein, and proximal vein (9 points) and their termination along the blade margin (9 points) were used. The landmarks form the framework for the rest of the points in the analysis, as they are homologous features found in every leaf. Landmarks are indicated as orange dots in **Fig. 1**. Between the landmarks, pseudo-landmarks were used to capture continuous curves, indicated in magenta in **Fig. 1**. The pseudo-landmarks were measured as a vector, an ordered set of spatial coordinate pixel values, with an origin and an end. The vectors are as follows: *m*, from the petiolar junction to the tip of the midvein; *d* from the petiolar junction to the tip of the distal vein; *p* from the petiolar junction to the tip of the proximal vein; *p1* (the petiolar vein), *d1,* and *m1* from the first secondary branch point of their respective primary veins to the termination of the vein at the margin; *p2, d2,* and *m2* from the second secondary branch point of their respective primary veins to the termination of the vein at the margin; *p3, d3,* and *m3* from the third secondary branch point of their respective primary veins to the termination of the vein at the margin; *pa*, *da, ma* along the margin from the beginning of their respective lobe to the termination of *p1*, *d1*, and *m1*, respectively; *pb, db, mb* from the termination of *pa*, *da,* and *ma*, respectively, to the termination of *p2*, *d2*, and *m2*, respectively; *pc*, *dc*, *mc* from the termination of *pb*, *db,* and *mb*, respectively, to the termination of *p3*, *d3*, and *m3*, respectively; *pd, dd, md* from the termination of *pc*, *dc*, and *mc*, respectively, to the tips of the proximal, distal, and midveins, respectively; *ps* and *ds* from the tip of the proximal and distal veins, respectively, to the midpoint of the proximal and distal sinus, respectively. The vectors are visualized as arrows in **Fig. 1**.

Vectors were traced by hand in ImageJ using the segmented line tool with fitted splines (Abràmoff et al., 2004). The set of *x* and *y* coordinates for each vector were saved as individual tab-delimited.txt files named by 1) the photo ID of the leaf indicating the vineyard position of the vine it was collected from, 2) an enumerating value 1 through 4 specifying which of four leaves for the variety the data corresponds to, and 3) which vector the data file represents. These files, the raw data, are available at the following link: https://github.com/DanChitwood/grapevine_ampelometry/tree/master/0_visual_check/ampel_ometry_data. Tracing all data for a single leaf took approximately 15 minutes. Because the data was traced by hand, it was important to visually verify its accuracy. Analyses in Python were undertaken using NumPy (Oliphant, 2006), pandas (McKinney, 2010), and Matplotlib (Hunter, 2007) to plot the data on the actual photo. The code for plotting vectors onto the original photo can be found here: https://github.com/DanChitwood/grapevine_ampelometry/blob/master/0_visual_check/ampel_ometry_visual_check.ipynb. The visual checks for each of the 240 leaves analyzed in this study can be found here: https://github.com/DanChitwood/grapevine_ampelometry/tree/master/0_visual_check/output_visual_check

### Interpolation and Procrustes analysis

Once data for all 240 leaves were collected, an appropriate number of points to interpolate for each vector was determined. Procrustes analysis requires corresponding points in every sample. For the 24 homologous landmarks, this condition is already met, but for the pseudolandmarks, an equal number of equidistant points for each vector must be calculated. A function to retrieve the overall distance of a vector path was created using the numpy.ediff1d function (consecutive differences between elements of an array) to calculate Euclidean distance and the numpy.cumsum function (cumulative sum of an array) to calculate the cumulative distance. For each vector, its total sum distance across all leaves was calculated, as well as the overall distance for all vectors for all leaves. The total number of landmarks + pseudolandmarks apportioned to a vector was based on its relative total distance. The total number of landmarks was chosen at 6,000. This was an arbitrary decision to select a number as high as possible so that pseudo-landmarks were saturating (creating continuous curves and capturing local details, such as serration shape) but still low enough that computationally intensive Procrustes analyses were feasible on a laptop computer. Due to rounding, the final number of landmarks was 5,999, assigned to vectors as follows: https://github.com/DanChitwood/grapevine_ampelometry/blob/master/1_interpolation/outp_ut_number_of_points.txt. With assigned numbers of points to every vector, interpolation was used to calculate equidistant pseudo-landmarks. A function was created using the scipy (Virtanen et al., 2020) interp1d function to interpolate the correct number of equidistance points for each vector. The code used to interpolate points is here: https://github.com/DanChitwood/grapevine_ampelometry/blob/master/1_interpolation/ampelometry_interpolation.ipynb. The interpolated points can be found here: https://github.com/DanChitwood/grapevine_ampelometry/blob/master/1_interpolation/output_interpolated_points.txt

With corresponding points between all leaves, a Procrustes analysis could be performed. Generalized Procrustes Analysis (GPA) minimizes distances between corresponding points through translation, rotation, scaling, and reflection to an arbitrarily selected reference shape. The resulting mean shape for the superimposed points is calculated and becomes the new reference if the Procrustes distance to the reference does not meet a minimum threshold (Gower et al., 1975). GPA was performed using the procGPA() function from the package “shapes” (Dryden and Mardia, 2016) in R (R Core Team, 2019). GPA was first performed for the four leaves for each variety producing mean shapes and superimposed Procrustes coordinates. The Procrustes mean shape and coordinates were used for plotting. The procdist() function from “shapes” was used to calculate the Procrustes distance between each pair of mean shapes and the results saved as a pairwise distance matrix. The hclust() function in R using the “mcquitty” method was used to hierarchically cluster varieties based on the pairwise distance matrix and overall morphological similarity. The code for performing a Procrustes analysis for each variety and outputs can be found here: https://github.com/DanChitwood/grapevine_ampelometry/tree/master/2_procrustes_by_variety

A GPA was also performed for all 240 leaves. The outputs include an overall Procrustes mean shape, super-imposed Procrustes coordinates for all leaves, and eigenvalues and eigenleaves from a PCA. The superimposed Procrustes coordinates of all leaves and the mean shape were plotted together. The code for the Procrustes analysis for all 240 leaves and the outputs can be found here: https://github.com/DanChitwood/grapevine_ampelometry/tree/master/3_overall_procrustes

### Data analysis

To calculate allometry for each line segment, distances between all points were converted to cm using the pixel to cm scale measured for each leaf. The lm() function in R was used to model the natural log of the distance from each point to the next as a function of the natural log of the overall distance for each leaf. The slopes and residuals were saved. Slope values for each point were projected onto the Procrustes mean leaf and visualized using ggplot2 (Wickham, 2016). The standard deviation of the residuals for each point was also calculated and plotted onto the mean leaf.

To calculate the statistical contribution of each landmark to discriminating leaves by variety, the Euclidean distance of each point to the corresponding point of the mean leaf was calculated. The distance of each point to the mean was then modeled as a function of variety using the kruskal.test() function. The test statistic and p-value were saved. The p-value was multiple test adjusted using the Benjamini-Hochberg method and plotted on the mean leaf.

To predict variety from leaf shape, datasets were first converted into orthogonal components using Principal Component Analysis (PCA) with the prcomp() function in R. Transformation into orthogonal variables was a necessity before proceeding with Linear Discriminant Analysis (LDA) to avoid collinearity (a problem with the saturating number of pseudo-landmarks with similar values used in this study). LDA was performed using the lda() function with the “MASS” package (Venables and Ripley, 2002). The cross-validated “leave-one-out” approach was used to predict the variety of each leaf using CV = TRUE. The confusionMatrix() function from the package “caret” (Kuhn, 2008) was used to calculate overall classifier statistics and estimates of accuracy from the resulting LDA model.

## RESULTS

### Morphological similarity, comparison to the results of Pierre Galet, and average leaf shapes

Using the pairwise Procrustes distance matrix of the overall morphological similarity of the average leaf of every variety to the other, a hierarchical clustering was performed to find groups of varieties with similar leaf shapes (**Fig. 2**). Because the clustering reflects the minimization of the distance of 5,999 points for each variety to the other, it is difficult to interpret which features of the leaf most strongly contribute to a leaf resembling another. To help understand which shape attributes of the leaf contributed to the clustering signal, the measurements of Pierre Galet for each variety were analyzed. The 60 varieties chosen for this study are included in Galet’s *A Practical Ampelography* (Galet, 1979; 1985). Each variety has values for the “Galet formula”, a method that measures the relative lengths of veins and their angles (**Fig. 1**). The values A, B, and C measure the relative ratio of the lengths of L2, L3, and L4, respectively, to the L1. The variable *r* is the ratio of length to width. S’ and S are angles between the L1 and the L3 and L4, respectively. Su and In are the ratios of distances from the petiolar junction (0) to the superior and inferior sinuses, respectively, divided by the length of the L2 and L3, respectively. Ratios and angles are discretized into values 0-9 and can be measured using the Galet ruler and the Galet protractor. For ratios of primary veins A, B, and C, increasing values correspond to decreasing ratios. For length-to-width ratio *r*, increasing values correspond to increasing ratios. For angles S’ and S increasing values correspond to increasing angles, and for measures of sinus depth Su and In, increasing values correspond to deeper sinuses. Comparing Galet formula values to hierarchical clustering, the overwhelming correspondence between the two datasets is sinus depth (Su and In; **Fig. 2**). Excluding uniquely shaped varieties that cluster alone (Chasselas cioutat, Zinfandel/Primitivo, Gewürtztraminer, and Burger/Monbadon), two major groups of varieties arise. Group I leaves are deeply lobed and Group II leaves slightly lobed or entire.

**Figure 2:**
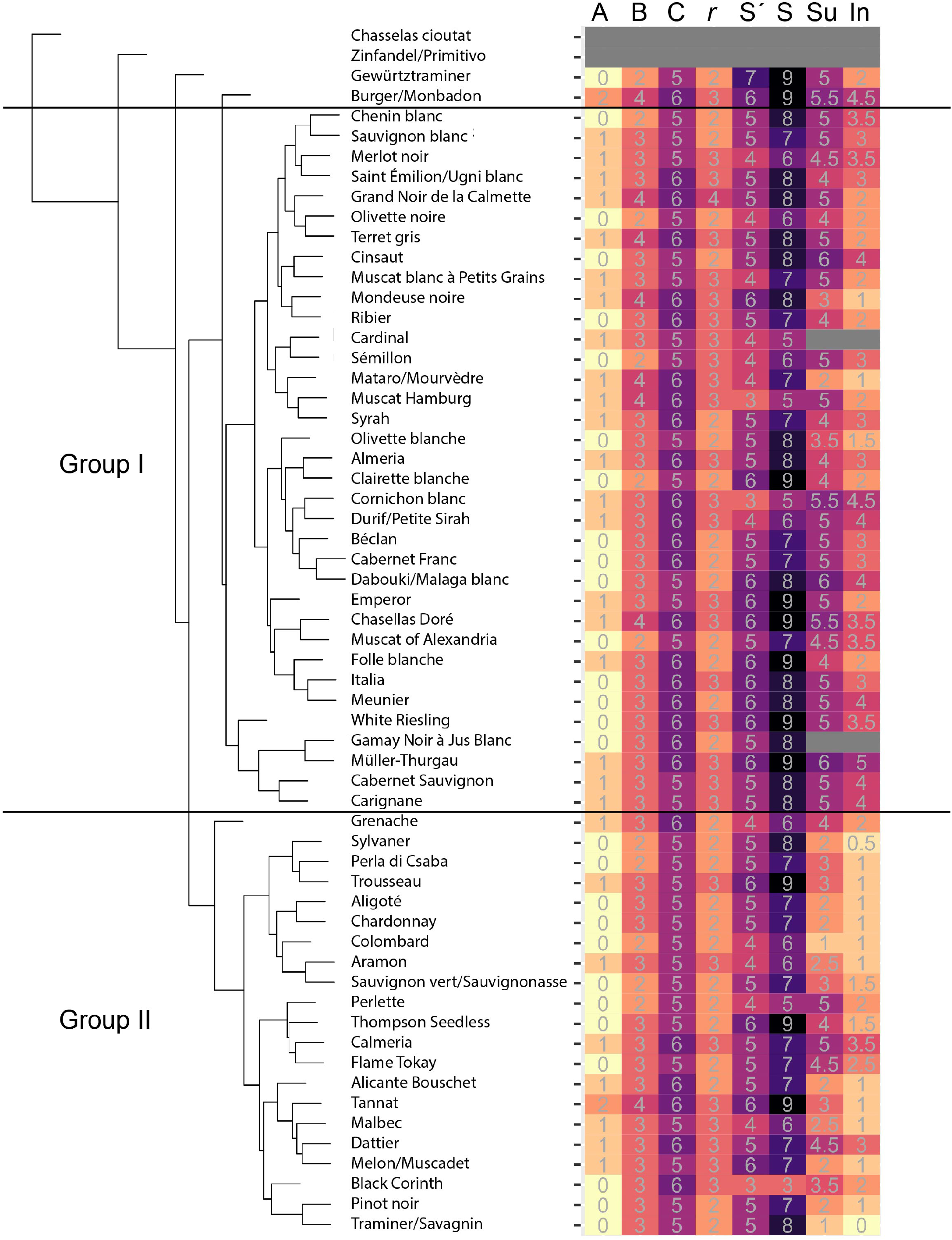
Clustering based on Procrustes distances and a comparison to Galet formula values. Hierarchical clustering based on a pairwise Procrustes distance matrix of the overall morphological similarity of averaged leaves for each variety is shown on the left. On the right, Galet formula values (colored from light to dark for low to high values) for A, B, C, *r*, S’, S, Su, and In, as defined in **Fig. 1**, are shown. Two Groups (I and II) with deep and slight lobing, respectively, are indicated.

One of the most impactful features of *A Practical Ampelography* (Galet, 1979; 1985) is Galet’s drawings. For each variety, Galet drew a representative leaf. While the Galet formula provided a means to quantify shape, the drawings capture the totality of information embedded in leaf shapes that we so easily take in with our eyes but defies measurement. The relationship of all angles comprising a leaf together, the curves of the primary and secondary veins, the shapes of the serrations, the shape of the petiolar sinus, and the overlap of lobes: these are features that impact the values of the Galet formula but are not fully captured by it. The drawings of Galet highlight the ampelographic features used to quantify grapevine leaves: namely, the veins and their relationship to the blade. By analyzing a saturating number of pseudo-landmarks, these intricate features of grapevine leaves have been quantitatively captured. To create a statistical version of Galet’s drawings, the 5,999 coordinate values for the four leaves for each variety were superimposed and the average leaf calculated. **Figs. 3–5** show the superimposed Procrustes coordinates for the four leaves for each variety (left), the average leaf (middle), and one example leaf with its coordinates overlaid. Such visualization combines the best attributes of landmark-based analyses and hand drawings: the calculation of an average leaf and the visualization of variance using superimposed Procrustes coordinates adds statistical rigor that drawings lack, while the use of a saturating number of pseudo-landmarks captures the continuous curves of veins and blade that a finite number of landmarks cannot. Leaves in **Figs. 3–5** are displayed in the order of their clustering in **Fig. 2**. At a glance, the deep lobing of Group I leaves in **Fig. 3** and **Fig. 4** can be compared to the more entire Group II leaves in **Fig. 5.**

**Figure 3:**
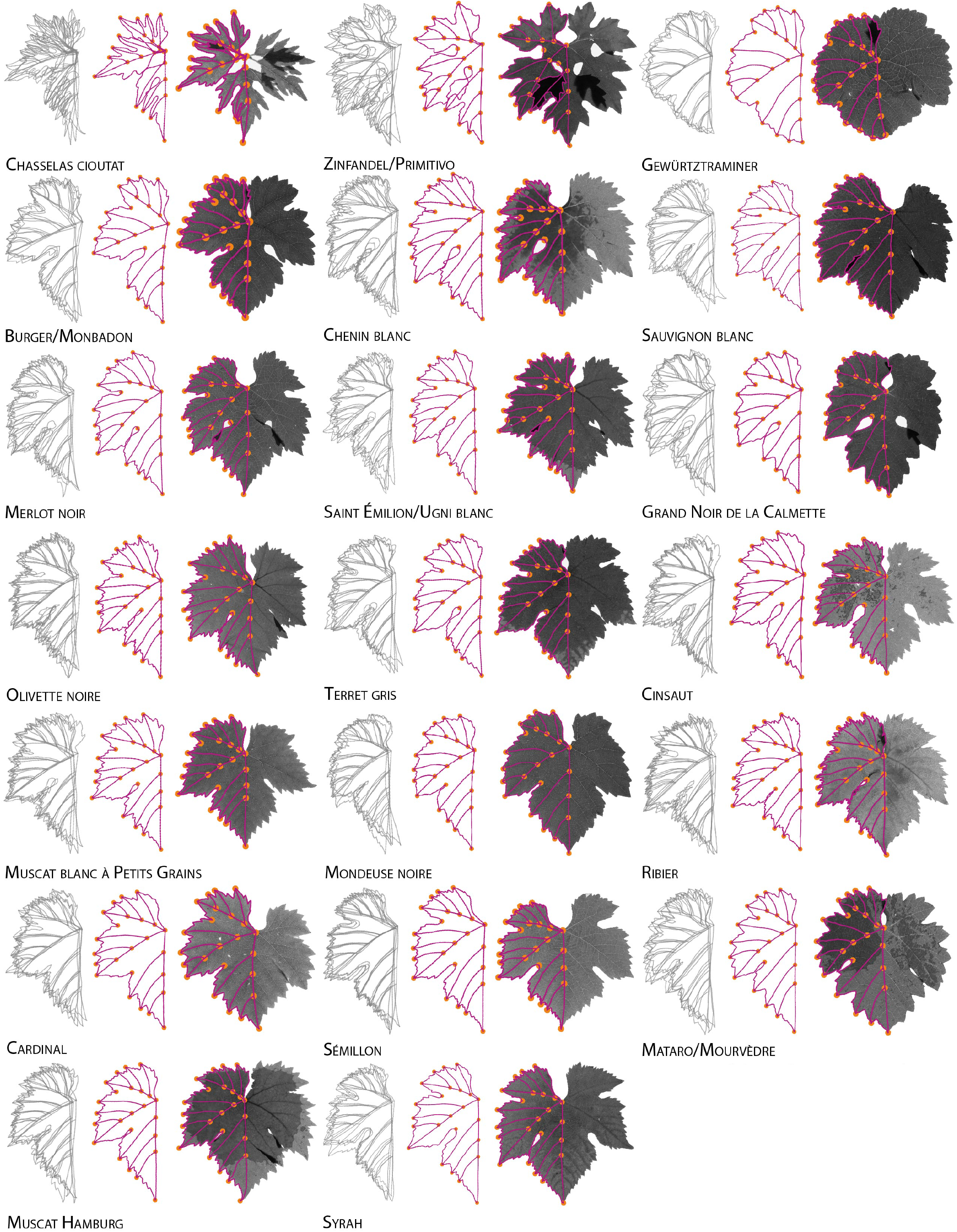
Leaf shapes by variety. Four leaves for each variety with superimposed Procrustes coordinates (left, gray), the mean leaf (middle, magenta and orange), and one example leaf overlaid with coordinates (right) are shown. Leaves are in the same order as presented in the hierarchical clustering in **Fig. 2**.

**Figure 4:**
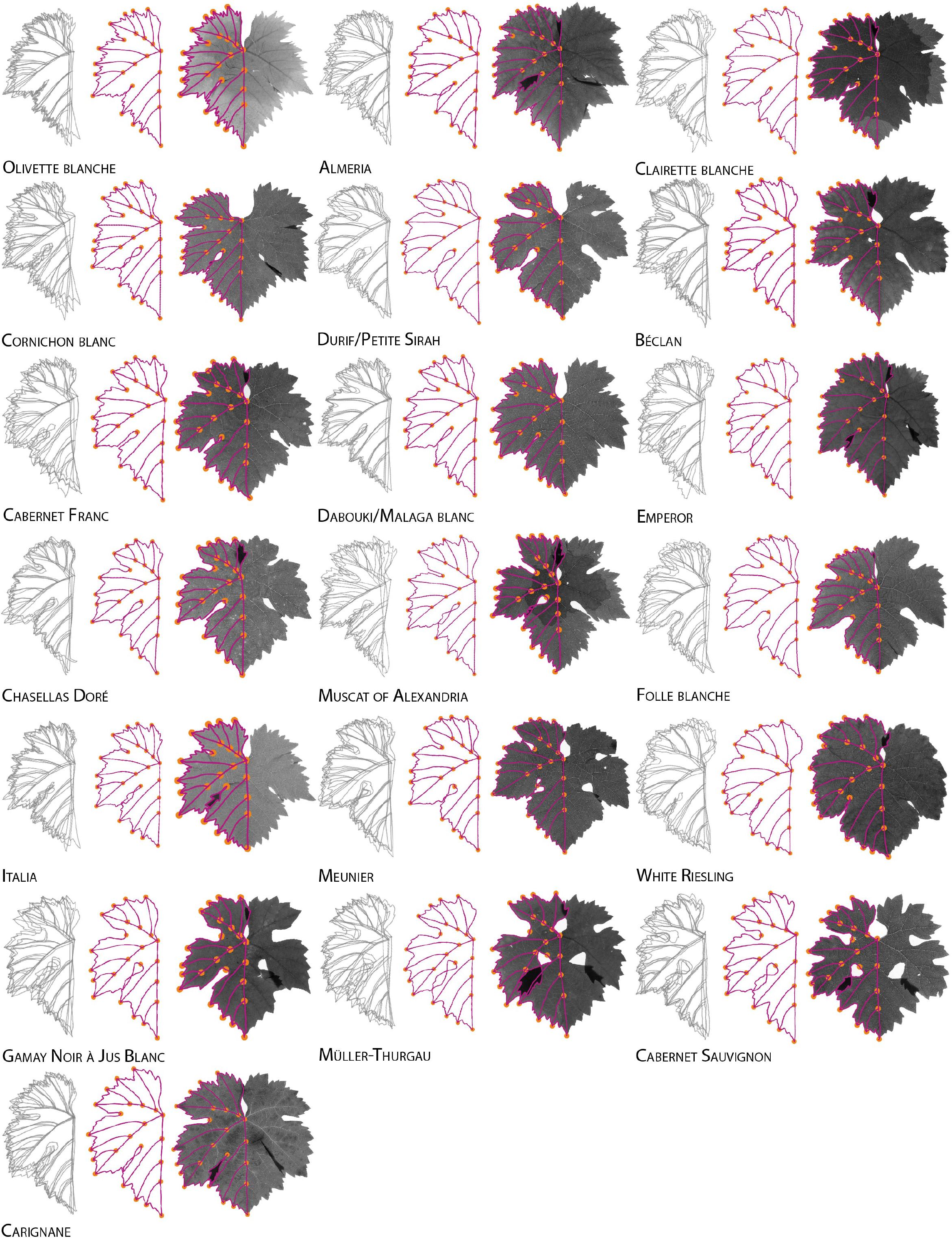
Leaf shapes by variety. Four leaves for each variety with superimposed Procrustes coordinates (left, gray), the mean leaf (middle, magenta and orange), and one example leaf overlaid with coordinates (right) are shown. Leaves are in the same order as presented in the hierarchical clustering in **Fig. 2**, continued from **Fig. 3**.

### Allometry and the ability of each coordinate to discriminate varieties

In order to analyze the contributions of individual coordinates to global features of the leaf and variability among varieties, a Generalized Procrustes Analysis (GPA) was calculated for all 240 leaves. All superimposed coordinates were overlaid on the overall average leaf (**Fig. 6A**). The mean leaf was subsequently used to project attributes of individual coordinates. Allometry (the differential growth of features in relation to organ size) was analyzed for each landmark. Previously, we demonstrated strongly linear relationships between the natural log of primary vein area vs. the natural log of blade area: smaller leaves have a higher vein-to-blade area ratio than larger leaves (Chitwood et al., 2016b). To determine the allometric relationships for the coordinates used in this study, the natural log of the Euclidean distance of each point to the next was regressed against the overall Euclidean distance of all veins and blades. The slope for each coordinate was recorded and plotted on the mean leaf (**Fig. 6B**). The distal/superior sinus had the largest slope values, demonstrating that relative to other segments of the leaf, the invagination of this region in deeply lobed varieties takes up a larger proportion of the overall leaf. The proximal side of the proximal/inferior sinus also has relatively high slope values. Although slight, for the mid and distal lobes, the slope is less at the tip and increases incrementally along the blade towards the base. This is consistent with the distal regions of the leaf and lobes initiating and developing before the proximal regions (Jones et al., 2013). To determine if there was a relationship between higher slope values and variability, the standard deviation of the residuals of the allometric regression were projected onto the mean leaf (**Fig. 6C**). Again, the distal/superior sinus and the proximal side of the proximal/inferior sinus had the highest variability. Together, the results show that the invagination of the sinuses, especially the distal/superior sinus, across varieties is the most malleable part of the grapevine leaf contributing to variation in leaf shape.

To determine the ability of different coordinates to discriminate varieties, a Kruskal-Wallis test was used. The Euclidean distance of each coordinate to the mean leaf was calculated and modeled as a function of variety. If the replicated leaves of one or more varieties consistently varies from the mean leaf, the Kruskal-Wallis test statistic will be responsive. After multiple test adjustment, coordinates in the distal/superior sinus were found to be the most significant, especially the points in the middle of the sinus pocket (**Fig. 6D**). The proximal/inferior sinus did not show similar variation between varieties, demonstrating that a strong allometric relationship (**Fig. 6B**) is not necessarily indicative of variability. The mid lobe showed the least significant variation between varieties. Not only is the distal sinus an allometrically sensitive region of the leaf, but it is one of the strongest indicators of variety, consistent with the depth of sinus lobing differentiating the two main morphological groupings of grapevine leaves (**Fig. 2**).

### Comparing the ability of different morphometric methods to predict variety

The morphometric methods presented so far rely on two embedded features: 24 homologous landmarks found in every grapevine leaf, and a set of 5,999 equidistant pseudo-landmarks that capture finer features, such as curves and serrations. Pierre Galet proposed a separate method of quantification, focusing on the ratios of lengths of lobes and relative angles between them (**Fig. 1**). He even published idealized values for each variety (**Fig. 2**) that could be compared with real world measurements by viticulturists using the Galet ruler and protractor. Without replication, there is no way to compare the methods of Galet to other morphometric techniques. In order to approximate the focus of Galet’s methodology on length ratios and angles, while preserving the continuous measurement of local features (such as curves and serrations) enabled by using a saturating number of landmarks, a ratio/angular transformation of the data was developed. For each coordinate 1) the ratio of its distance from the petiolar junction divided by the length of the midvein and 2) its angle from the midvein was calculated. Plotting the ratio of the distance from the petiolar junction against angle, features of the leaf are still apparent (**Fig. 6E**). The mid lobe, as the point of comparison, lacks variability. But the farther from the mid lobe points lie, the more variation is observed. This is in part because of variation in the primary vein angles, which was a focus of the methodology of Galet and Ravaz. The petiolar vein, in particular, shows a large amount of angular variation relative to the midvein, verifying the longterm focus of ampelographers on the petiolar sinus as a source of identifying information between varieties.

With replication for three different methods (only the 24 landmarks, the Galet-inspired transformation to ratios and angles, and all 5,999 Procrustes-adjusted coordinates) the ability to predict variety from shape information can be compared. A Principal Component Analysis (PCA) was performed on all three datasets to reduce information into orthogonal components. This step was necessary to avoid the collinearity of points that are, by definition, colinear. A Linear Discriminant Analysis (LDA) was performed on increasing number of PCs using a crossvalidated approach and the overall accuracy recorded. Each method peaked in accuracy and then diminished (**Fig. 6F**). For the only landmark method the peak in accuracy was at 27 PCs, for the Galet-inspired method 42 PCs, and for the all Procrustes coordinate method at 54 PCs. The amount of variation in the higher number PCs is miniscule (**Fig. 6G**) yet still contributed to increases in model accuracy. This demonstrates that especially for the Galet-inspired and all Procrustes methods, that fine details captured by higher order PCs still contain relevant information to discriminate between varieties. Plotting out the prediction from each dataset as a confusion matrix, especially for the only landmark dataset with lower accuracy, leaves tend to be most often confused within Groups I and II (**Fig. 7A**). The increased accuracy of the Galet and all Procrustes methods is expected given the increased amount of information that is captured using a saturating number of pseudo-landmarks (**Fig. 7B-C**). The overall accuracy of the only landmarks method was estimated at 0.454 (95% confidence interval 0.390 to 0.519, p-value = 5.70 x 10^-125^), whereas the accuracy of the Galet method at 0.579 (95% confidence interval 0.514 to 0.642, p-value = 5.72 x 10^-179^) and the all Procrustes method at 0.629 (95% confidence interval 0.565 to 0.690, p-value = 2.04 x 10^-202^) shows that saturating numbers of landmarks— regardless of method—contributes to increased accuracy in predicting variety.

## DISCUSSION

Leaf shape has historical importance in grapevines. Had genotyping existed in the late 1800s, new rootstock varieties to combat phylloxera in Europe and the North American *Vitis* spp. parents from which they are derived would have been identified molecularly. However, molecular biology did not exist yet. To verify rootstock identity and enforce appellation laws, the earliest of ampelographers, Goethe and Ravaz, turned to the angles and shapes of the petiolar sinus. Before the concept had existed, a relationship between genotype and phenotype, based on leaf morphology, was used to enforce law and regulate trade. Pierre Galet took the concept further, extending a framework for measuring the ratios of vein length and their angles to capture overall leaf morphology, as well as cataloging shape through hand-drawings, allowing readers to appreciate the beauty of grapevine leaf diversity and its constituent features at a glance. María-Carmen Martínez examined the features of leaves in even greater detail, allowing averaged leaves to be reconstructed at the level of individual teeth along the margins and providing inspiration for landmark-based methods. Using landmarks, genetic, developmental, and environmental effects on leaf shape have been measured. Yet, the high replication that a limited number of landmarks permits misses the exquisite features of veins and blade, while drawing-based methods that holistically capture the leaf have until this point been difficult to quantify.

Using a saturating number of pseudo-landmarks that capture continuous curves and intricate local features, powerful Procrustean-based methods can be used to measure leaf shape at a global level. A pairwise Procrustes distance matrix clusters leaves into two major categories: deeply lobed and more entire (**Fig. 2**). These categories correspond to Pierre Galet’s measurements of sinus depth, showing that this feature especially is diagnostic of variety, even when varieties are measured on different continents and decades later. Calculating the Procrustean mean shape is a way to summarize drawings quantifying underlying replication, preserving local and global features to represent the ideal leaf for each variety without having to pick any particular individual one as an example (**Figs. 3–5**). The distal/superior sinus contributes disproportionately to the variation in leaf shape that discriminates varieties, both through allometry and the conspicuous placement of the distal/superior sinus pocket (**Fig. 6A-D**). The focus of Galet on the ratios of vein lengths and relative angles can be used to transform continuous coordinates while preserving the overall morphology of leaves (**Fig. 6E**). Both the Galet-inspired transformation to ratios and angles and using all Procrustes-adjusted coordinates gives comparable overall accuracies of 0.579 and 0.629, respectively (**Fig. 6F-G**, **Fig. 7**). The much lower accuracy using only the 24 landmarks of 0.454 shows that the use of saturating pseudolandmarks (and less the framework within which they are analyzed) leads to higher prediction rates through capturing intricate features of the leaf. Using Procrustean pseudo-landmarks quantifies grapevine leaf shape to the same level of detail as drawings and provides a quantitative method to arrive at mean leaf shapes representing varieties that can be used within a predictive statistical framework.

**Figure 5:**
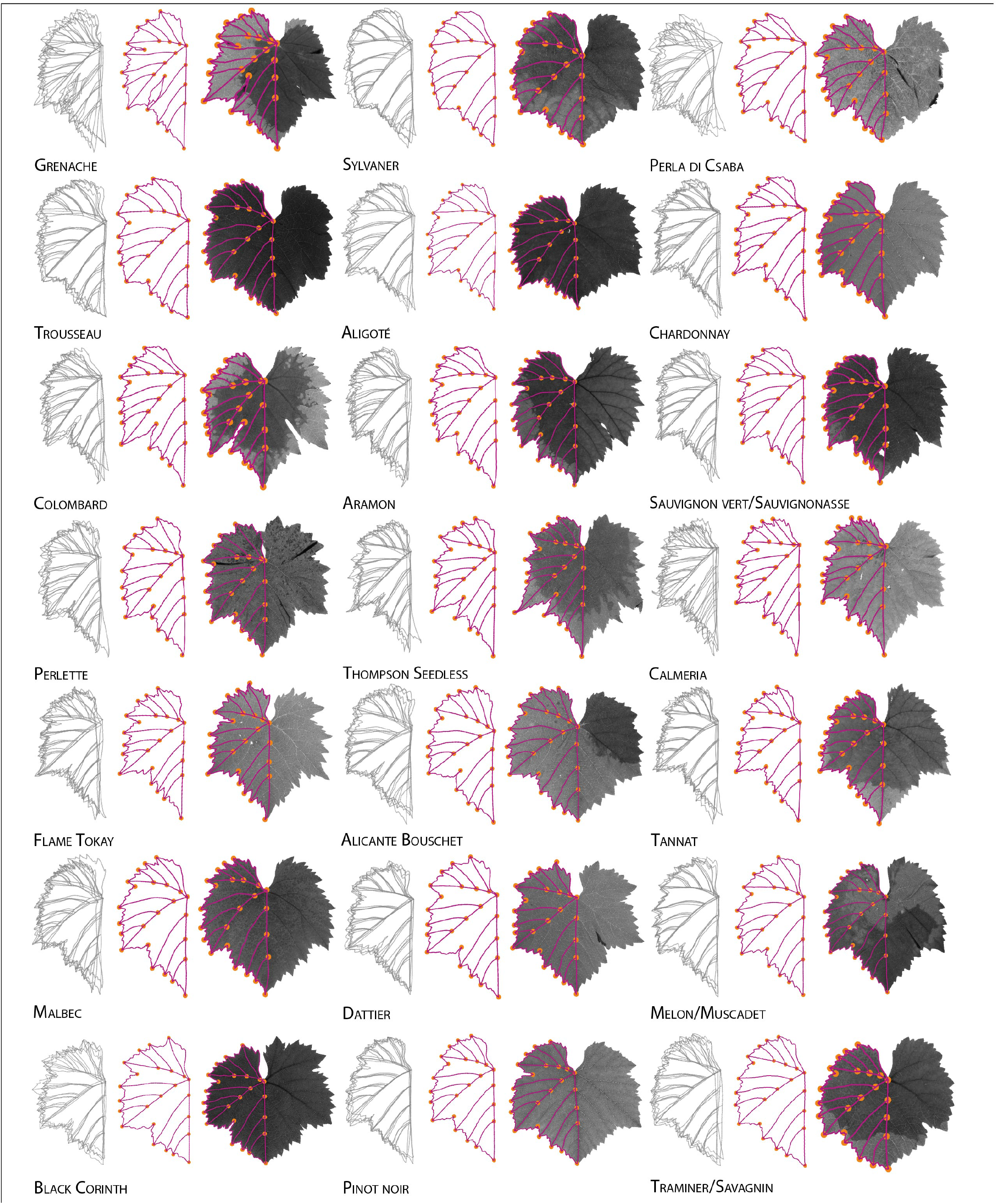
Leaf shapes by variety. Four leaves for each variety with superimposed Procrustes coordinates (left, gray), the mean leaf (middle, magenta and orange), and one example leaf overlaid with coordinates (right) are shown. Leaves are in the same order as presented in the hierarchical clustering in **Fig. 2**, continued from **Fig. 4**.

**Figure 6:**
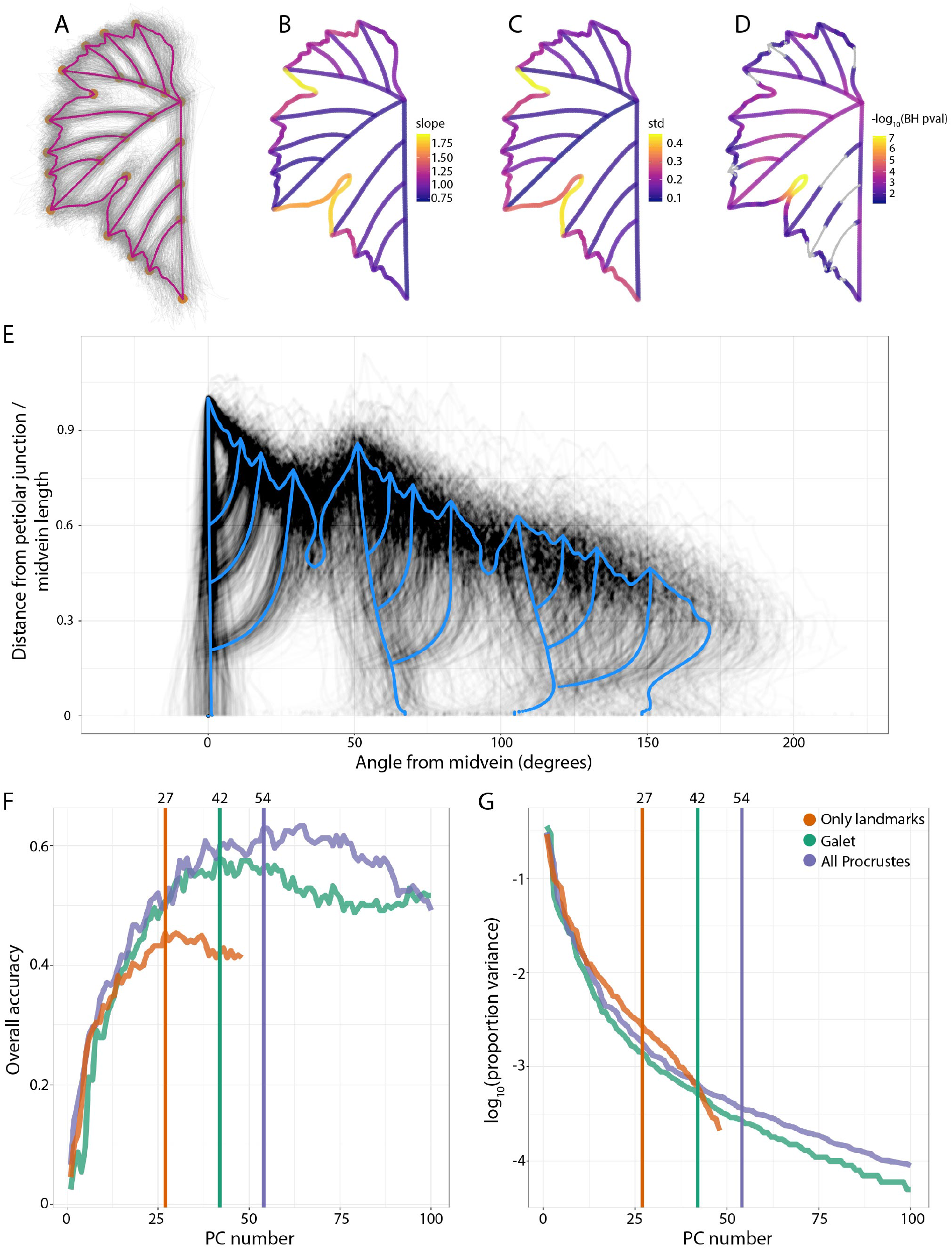
Allometry and variability between varieties and their prediction. **A)** Superimposed Procrustes coordinates for all leaves (gray) and overall mean leaf (magenta and orange). **B)** Allometric values for each coordinate projected onto the mean Procrustes leaf. Points are colored by slope of a fitted line for the natural log of the given point to the next divided by the natural log of the overall total distance of the leaf. **C)** Mean leaf with coordinates colored by the standard deviation of the residuals for each coordinate for the allometric relationship described in B. **D)** Mean leaf with coordinates colored by -log_10_ p-values (Benjamini-Hochberg multiple test adjusted) for a Kruskal-Wallis test for the Euclidean distances of each point to the mean leaf modeled by variety. Values failing to meet the adjusted significance value of p = 0.05 are shown in gray. **E)** A plot of the distance of each coordinate to the petiolar junction divided by the midvein length versus the angle of each point from the midvein (the angle defined by the tip of the midvein, the petiolar junction, and the point of interest). The mean leaf defined by angle and distance coordinates is shown in blue. **F)** Three morphometric methods are compared: only landmarks (24 landmark values, orange), the Galet-inspired method (angle and distance transformation, teal), and all Procrustes points (the 5,999 landmarks + pseudolandmarks, lavender). The overall accuracy of predicting variety using the indicated number of PCs for each method is plotted. The number of PCs that yielded the maximum accuracy ultimately used for prediction is shown (27 for only landmarks, 42 for Galet, and 54 for all Procrustes). **G)** The -log_10_ value of the proportion of variance explained by each of the PCs for each of methods is shown. Again, the number of PCs used for prediction that yielded the maximum accuracy is indicated.

**Figure 7:**
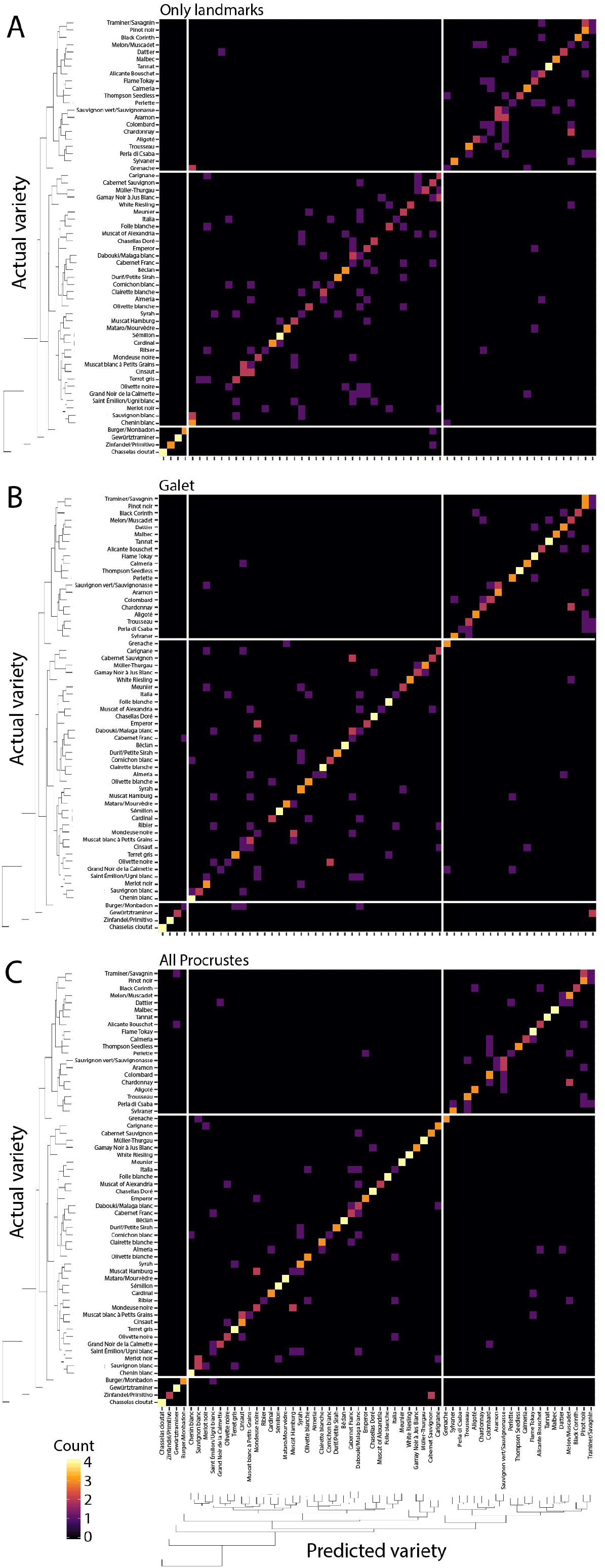
Predicting variety from shape. Confusion matrices showing the accuracy of prediction for leaves by variety for the **A)** only landmark (24 landmark values), **B)** Galet-inspired (angle and distance transformation), and **C)** all Procrustes (5,999 landmarks + pseudo-landmarks) methods. For each confusion matrix, varieties are arranged by clustering based on overall morphological similarity in **Fig. 2**. Group I and II varieties with deeply and slightly lobing leaves (respectively) are separated by white lines. The number of leaves assigned, zero to four, is indicated by color (dark to light).

## ACKNOWLEDGEMENTS

The author is forever grateful to Pierre Galet (1921 – 2019), whose work and vision has provided boundless scientific and artistic inspiration.

